# Global relationships between crop diversity and nutritional stability

**DOI:** 10.1101/2020.07.29.227520

**Authors:** Charlie C. Nicholson, Benjamin F. Emery, Meredith T. Niles

## Abstract

Nutritional stability – a food system’s capacity to provide sufficient nutrients despite disturbance – is a critical feature of sustainable agriculture, especially in light of ongoing climate change. Yet, measuring nutritional stability has proven challenging. Addressing this challenge will help identify resilient food systems, detect shortcomings in nutrient availability, and evaluate if stability-focused interventions actually work. We develop a novel approach that uses 55 years of crop data across 184 countries to assemble over 22,000 bipartite crop-nutrient networks. We then quantify the tolerance of these networks to disturbance simulated via sequential crop loss (Fig. 1) and evaluate patterns of crop diversity and nutritional stability across countries, over time and between crop supply scenarios (imports versus in country production). We observe a positive, saturating relationship between crop diversity and nutritional stability across countries; however there is substantial variability between countries over time. Next, despite crop diversity gains since 1961, nutritional stability has remained stagnant or decreased in all regions except Asia. A decline in the average number of nutritional links per network (range: -3 to -18% across regions) and the aforementioned saturating relationship explain this counter-intuitive finding. Finally, we find that imports increase crop diversity and improve or sustain stability, indicating that nutrient availability is market exposed in many countries, particularly developing states. Although applied globally, our approach is applicable across levels of organization, from household intake to sub-national production, and provides a way forward for understanding the contributions of crop diversity to the stability of nutrients available for human consumption.

## Introduction

Market volatility, land degradation, pests, and climate change make it increasingly difficult for agriculture to sustain a growing and healthy human population. In the face of these challenges, longstanding global-and national-scale policies seek to improve food security by increasing national production, leveraging international trade to counteract crop failures and supporting technological innovation (e.g. precision irrigation, drought resistant cultivars). These policies typically evaluate outcomes in terms of total yield or food calories, however expanding commitments to nutrition-sensitive agriculture (2, 3) has recognized that calories does not equate to food security. There is now increasing focus on nutritional diversity, including diet diversity and the diversity of nutrients needed to sustain a balanced diet and lead an active, healthy life (4).

With a growing focus on nutritional diversity, crop diversification is seen as a promising strategy to improve dietary diversity and nutritional status (5, 6). Nonetheless, there is mixed evidence linking crop diversity to nutritional outcomes (7), leading to calls for multilevel and systemic measurement approaches that can evaluate the influence of agrobiodiversity on the provision of nutrients at different scales (e.g., village, region, and national) (8–10). Potential nutrient adequacy (11), one advancement toward this end, incorporates into a single metric the fraction of a population potentially nourished for all nutrients by all crops. However, neither this, nor any other approach, measures the ability of agriculture to produce nutritious food through space and time in the face of chronic disturbance and acute shocks (12).

Quantifying this nutritional stability – the capacity of a food system to provide sufficient nutrients despite disturbance – can aid effective coordination and implementation of resilience planning for nutrition-sensitive agriculture. Target 3 of the Sustainable Development Goals stresses connecting crop diversity, resilient farming systems, and nutritious diets as essential components of food security (13). There are plausible mechanisms and demonstrated evidence for crop diversity contributing to human nutrition through the addition of key crops belonging to distinct nutritional functional groups (14–16). Yet, the fragility of this diversity-function relationship is unknown because a straightforward method to measure nutritional stability does not exist (1).

Here we develop a novel approach that measures the extent to which crop diversity underpins nutritional stability over space and time. Our analytical framework (**Fig. 1**) links crops and their constituent nutrients into a bipartite network and then quantifies the effect on nutrient availability in a given country when crops are removed from the network. This approach has characterized network tolerance in diverse fields, from information systems (17) to ecological networks (18), but has never been applied to better understand the stability of nutrient availability in food systems. As with most biodiverse communities, species removal order can structure loss of system function (19, 20). We therefore generate a generalized ‘attack tolerance curve’ (21) via permutation of removal sequence and derive our nutritional stability metric (*R*_*N*_) as the area under this curve (**Fig1d, e**; see Methods). A network with more nutrient dense crops will have more nutritional links (i.e. higher degree) and thus be more robust to crop loss.

**Figure 1.**
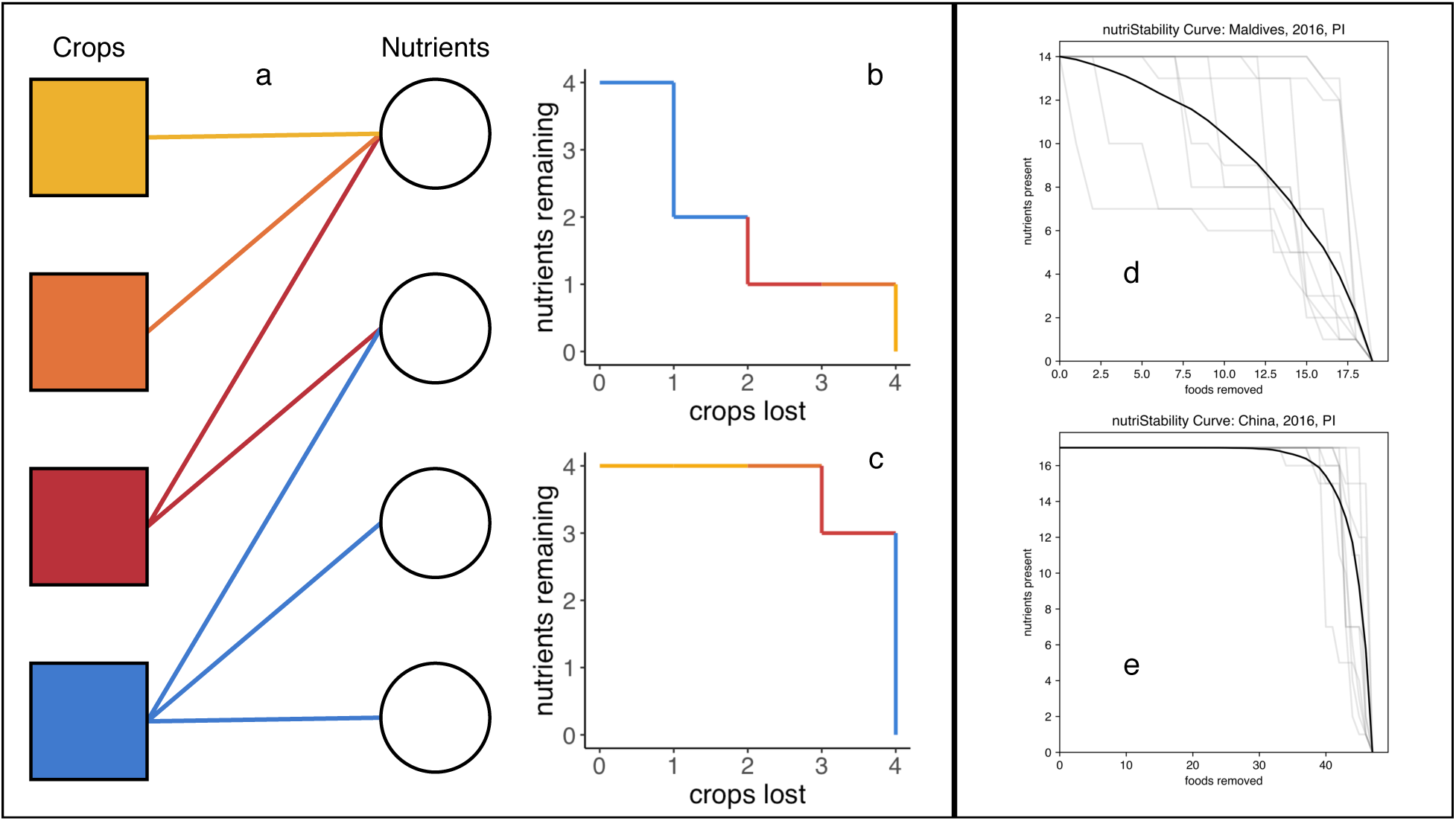
A new approach for understanding nutritional stability. The robustness of networks, such as a crop-nutrient network (a), can be gauged through assembling attack tolerance curves (b & c), whereby as nodes (i.e. crops) are eliminated constituent nutrients are lost. Of course, removal order matters (b vs c), therefore permuted combinations of crop loss generate average loss curves (see Methods). We use the area under this curve as our measure of nutritional stability (*R*_*N*_). Crop-nutrient networks with few redundant connections are susceptible to crop removal and can experience rapid nutrient loss (e.g. Maldives; d), whereas countries with a diverse food supply are more robust (e.g. China; e). This unitless metric is generalizable across different levels of aggregation (e.g. individual, household, community, national food system).

We apply this method to over 22,000 crop-nutrient networks assembled using 55 years of FAO food balance data for 184 countries and a global nutrient composition database (22). We evaluate patterns of nutritional stability – and its relationship with crop diversity and nutrient availability – across countries, over time and between two supply scenarios: food derived from production (P) and food from production and imports (PI). We then ask: (1) What is the relationship between crop diversity and nutritional stability? (2) How has crop diversity and nutritional stability changed over time in countries and regions? (3) What regional patterns underpin differences in nutritional stability?

## Results

Crop diversity begets nutritional stability with a non-linear relationship that has regional variability (**Fig. 2**). Across countries, nutritional stability increases with crop richness at significantly (p< 0.05) different rates between regions, generally achieving a threshold at which additional crops do not provide significant improvements in nutritional stability. (**Fig. 2, Fig. S1; Table S1**.). Across regions, gains in nutritional stability generally slow after crop-nutrient networks contain between 10 to 15 unique crops. Additional crops, beyond those present in those regions, do not provide significantly greater gain in nutrients available in those regions. This suggests there is a threshold for the extent to which adding more crop diversity into a region provides significant nutrient availability gains.

**Figure 2.**
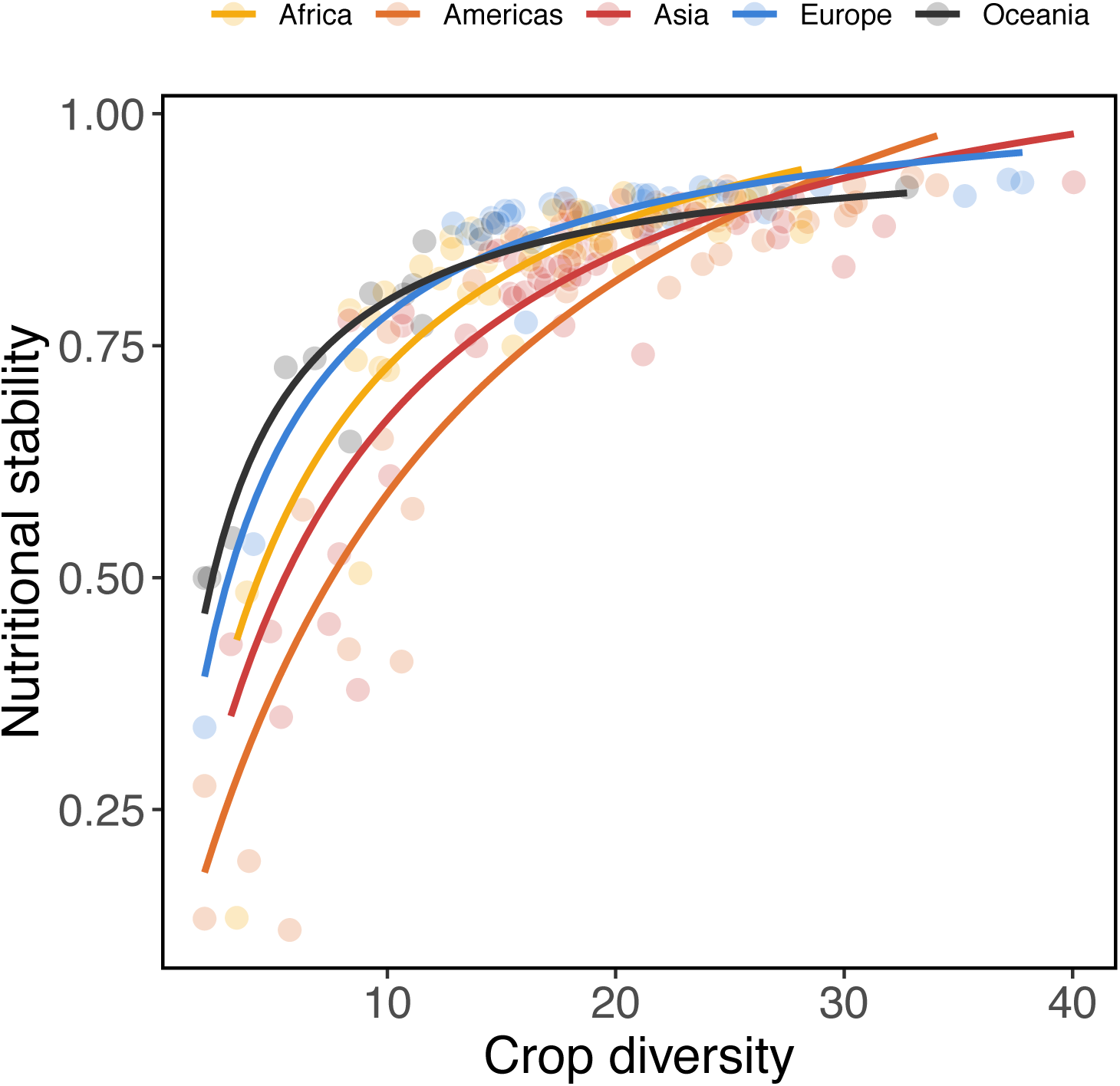
Nutritional stability increased non-linearly with crop diversity. Individual curve’s doubling-time parameter (β) differed significantly from each other (Fig. S1; Table S1), indicating region-dependent responses to crop diversity. Each point is a country average across all years.

We find that crop diversity increased since 1961 for all regions except in Oceania (**Fig. 3A, Table S2**). Importantly, there is significant variability across the remaining four regions, including the role of imports in driving crop diversity change over time. For example, advances are pronounced in Asia and Europe, where imported crop diversity increased by 43% & 35% between 1961 and 2016, respectively. Europe also has the largest gap between imports and production, with the majority of crop diversity increases driven by imports. In other words, increases in crop diversity at the country level are not necessarily (and frequently not) the result of increased diversity of production within a country.

**Figure 3.**
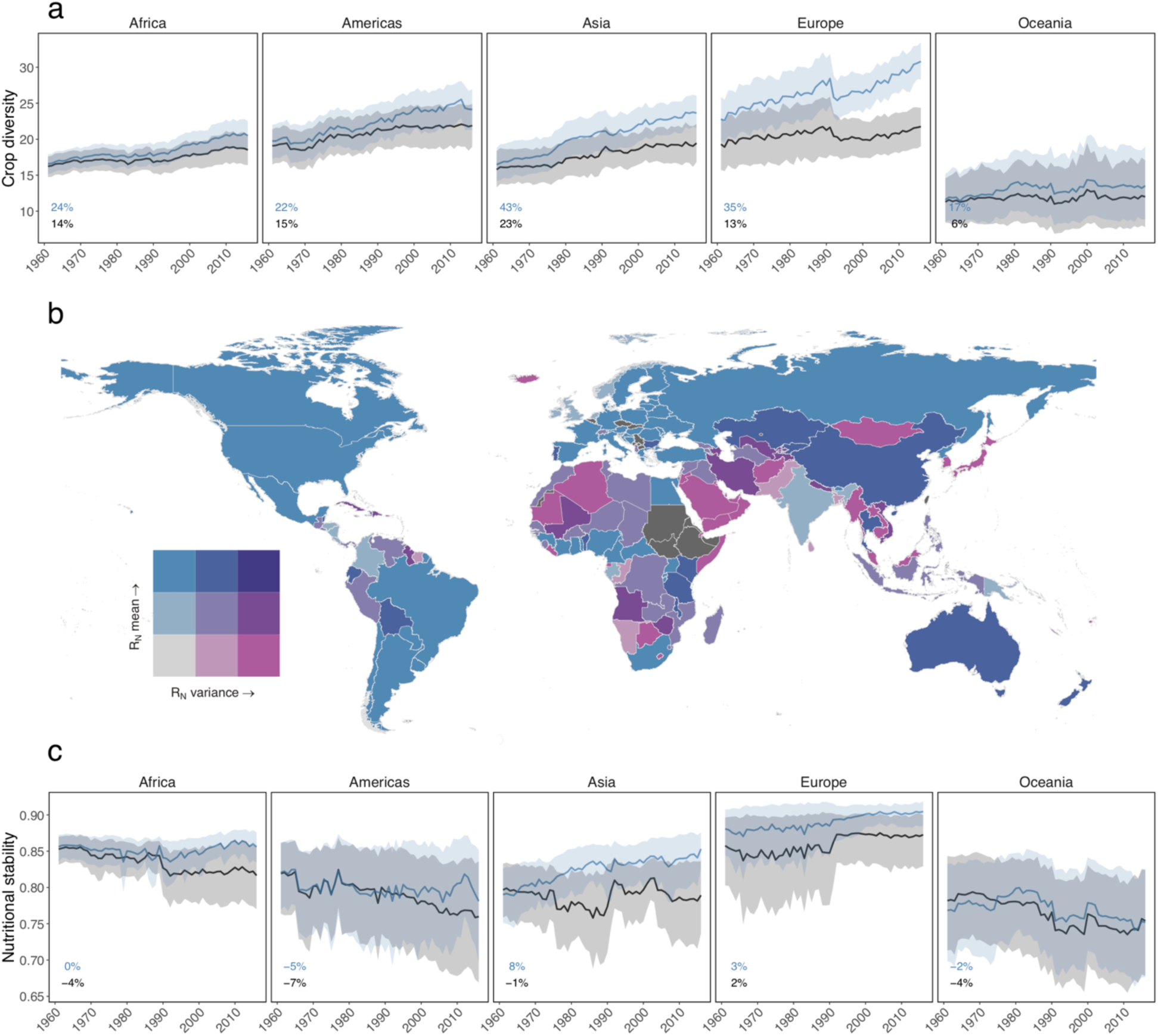
Global patterns and trends of crop diversity and nutrient stability. Crop diversity increased over the 1961-2016 period for production plus imports (blue) and production alone (black) scenarios (a). Nutritional stability (*R*_*N*_) varies over space (b) and through time (c). Bivariate choropleth maps depict R_N_ mean and variance across years. For example, light purple countries experienced low average nutritional stability and high year-to-year variation. Countries filled with dark gray lacked sufficient data. Trends in *R*_*N*_ (c) varied between region and depended on supply scenario (Table 1), with Africa and Asia exhibiting diverging responses for food supplies derived from production plus imports (blue) and production alone (black). Trend lines depict means ± 95% confidence intervals. The percentage change over the 55 year period for each regions and scenario is in the lower left of each panel.

We also find high variability in nutritional stability between countries over time (**Fig. 3B**). For example, countries such as the United States, Brazil, and much of Europe (light blue areas) had high stability with low variability in that stability (i.e. a small range of values measured as the variance in *R*_*N*_ over the period 1961 – 2016), meaning there was a consistently high stability of nutrients available in the country’s food supply. Conversely, other countries such as many in the Middle East, Southeast Asia, and Africa experienced low stability and large variance (light purple areas), indicating a highly variable and generally unstable supply of micronutrients available in these countries. Notably, small island developing states (SIDS), and low-income countries often experience low stability (**Fig. 4; Table S3**).

**Table 1.** Nutritional stability trends over time. Results are from region-specific linear mixed effects model with an interaction between supply scenario (production + imports v. production alone) and year as fixed-effects, country nested in scenario as random effects and an autoregressive correlation structure to account for temporal autocorrelation. Values are model coefficients with standard error in parentheses.

|  | Nutritional stability |  |  |  |  |
| --- | --- | --- | --- | --- | --- |
|  | Africa | Americas | Asia | Europe | Oceania |
| Supply Scenario | -1.360**<br>(0.412) | -1.391<br>(0.816) | -2.197*<br>(0.822) | -0.394<br>(0.306) | -0.452<br>(0.548) |
| Year | -0.0004**<br>(0.0001) | -0.001**<br>(0.0003) | -0.0003<br>(0.0003) | 0.0002*<br>(0.0001) | -0.001**<br>(0.0002) |
| Supply Scenario × Year | 0.001***<br>(0.0002) | 0.001<br>(0.0004) | 0.001**<br>(0.0004) | 0.0002<br>(0.0002) | 0.0002<br>(0.0003) |
| Observations | 5,546 | 4,217 | 4,577 | 3,076 | 1,628 |
| Log Likelihood | 13,648.020 | 9,471.530 | 9,456.794 | 7,189.631 | 4,561.369 |
| Note: | * p<0.05; ** p<0.01; *** p<0.001 |  |  |  |  |

**Figure 4.**
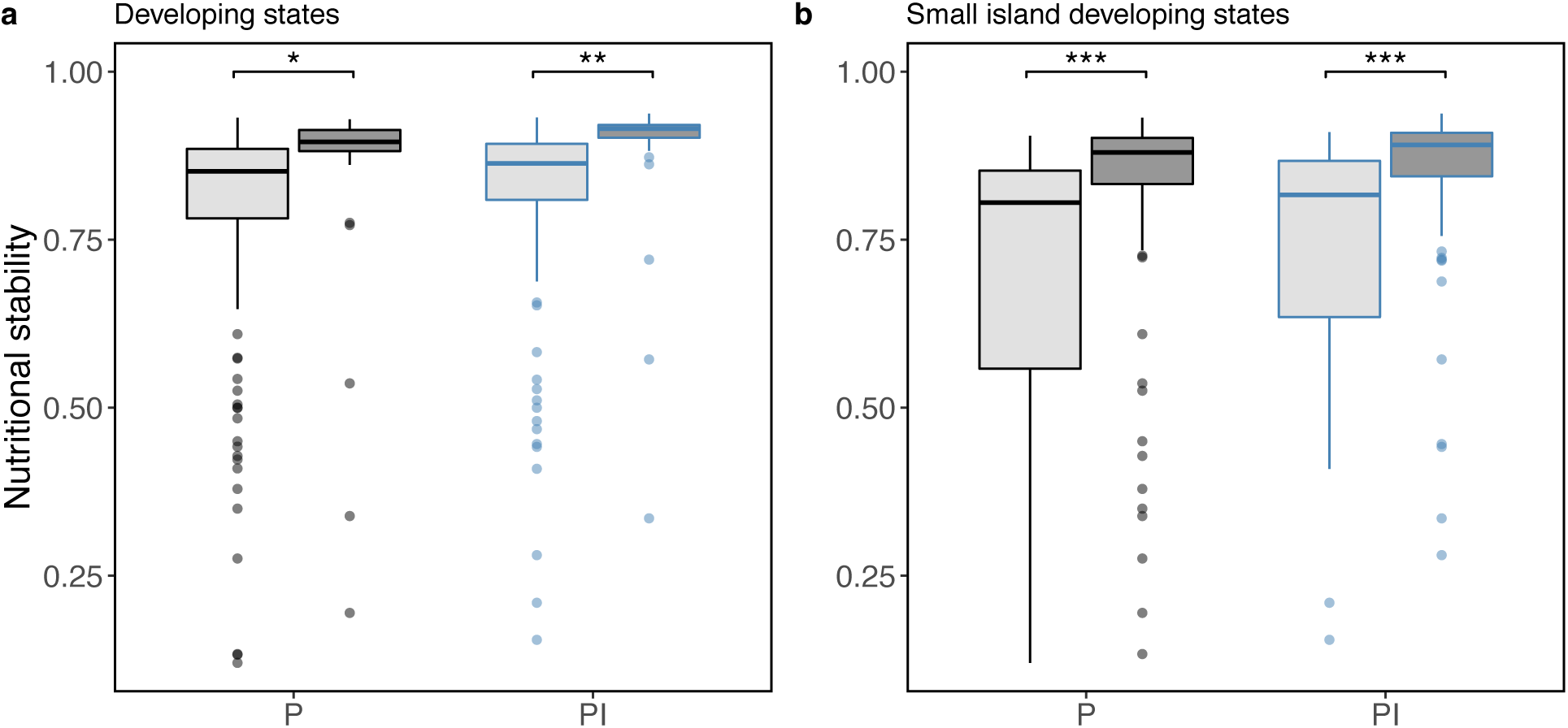
National factors related to nutritional stability. Developing states (a, light gray) and small island states (b, light gray) often experienced low nutritional stability. Each point is a country’s nutritional stability averaged across years (black: production alone; blue: production plus imports). Small island states are included in developing states. See Table S3 for statistical comparisons, * *P* < 0.05, ** *P* < 0.01, *** *P* < 0.001.

Despite finding that crop diversity is increasing, we find that nutritional stability has remained stagnant or even decreased in all regions except Asia (**Fig. 3C, Table 1**). Furthermore, across all regions, any gain in nutritional stability at the regional level over the time period are exclusively associated with a region’s supply of nutrients available from production and imports, not regional production alone. For example, while Asia experienced an 8% increase in nutritional stability, this was from an increase in imports over the time period; nutritional stability from production alone actually declined by 1%. Conversely, Africa experienced a 4% decrease in nutritional stability derived from reductions in crop diversity at the production level alone, while import-based nutritional stability did not change between 1961 and 2016. European nutritional stability was both relatively high (compared to other regions) and stable across the time period. In the Americas and Oceania, we find production-based nutritional stability decreased by 7% and 4% (respectively).

We present a seemingly counter-intuitive finding: for most regions crop diversity has increased yet nutritional stability is stagnant or decreasing. The saturating relationship between crop diversity and nutritional stability in part explains this, but changes in crop degree – the average number of nutrients provided by each crop in a network – clarify this incongruity further (**Fig. 5**). While crop diversity from production and imports increased for 72% of countries, 87% of these countries saw the average crop degree decrease (**Fig. 5**, upper left), and this is associated with a decrease in nutritional stability. In fact, crop degree has declined in all regions over the 55 year period, regardless of supply scenario (**Fig. S2, Table S4**). Stated succinctly, even though crop diversity has increased, there are diminishing returns on nutrient availability from adding more crops into a network, especially since the crops being added into networks appear to provide fewer links to nutrients not already in the food system.

**Figure 5.**
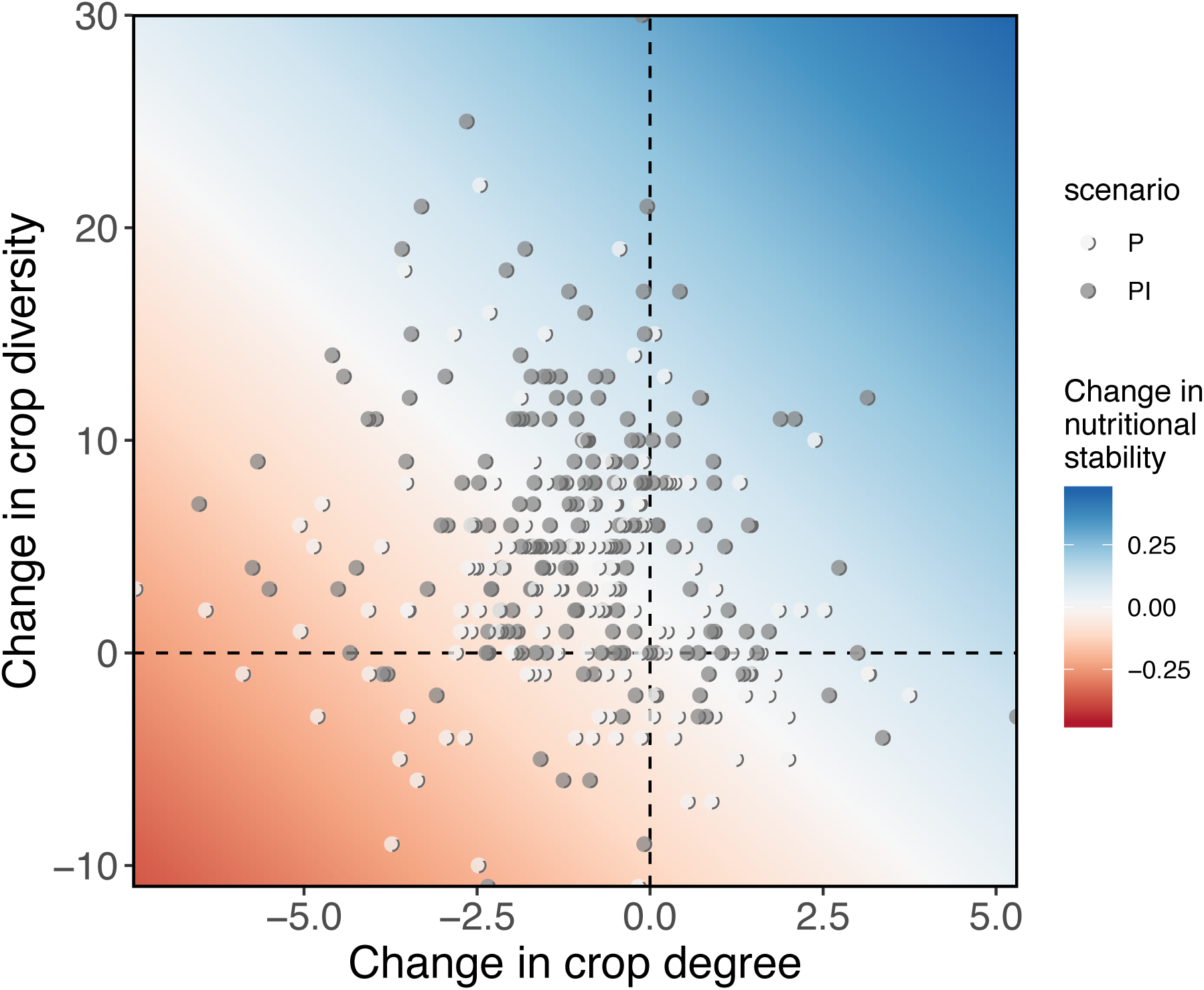
Country-level changes in crop diversity, degree and nutritional stability. Each point depicts the difference between a country’s last (2016) and first annual record of crop diversity (i.e., number of crops) and crop degree (i.e., average number of nutrients connected to each crop) for production (P; white circles) and production plus imports (PI; gray circles). The response surface projects the expected change in nutritional stability as predicted by simple linear regression (R_N_ ∼ diversity + degree).

## Discussion

Narrowing crop diversity in the world’s food supplies has been considered a potential threat to food security (23); however, there have been few empirical studies to link crop diversity to nutritional outcomes, especially beyond dietary intake at the household level (10). Here we develop a novel method to link crops to specific micronutrients using a network approach and assess the role of crop production and imports on nutritional stability outcomes in 184 countries between 1961 and 2016. Similar to other scholars (23, 24), we find that crop diversity has increased over time in all regions except Oceania, but that in many cases these gains are due to imports. Despite this increase in crop diversity, nutritional stability has remained stagnant or decreased in all regions except Asia, a trend largely attributed to our finding that recent crop diversity gains result in fewer new nutritional links in a given food system.

The general relationship between crop diversity and nutritional stability is contextualized by changes in crop degree, and explains why stability trends do not mirror diversification. Improving crop diversity will always increase the size of the crop-nutrient network, but stability depends on the number of links within this network. As in other diversity-stability relationships functional identity matters, and declines in crop degree could reflect shifts towards networks with less nutrient-rich crops. For example, production-based crop diversity in Senegal increased by 29%, while crop degree dropped by 19% as the composition of its food supply shifted from staples (e.g. millet, groundnuts, sweet potatoes) to include less nutrient dense crops (e.g. sugar cane, watermelon, cabbage). In light of on-going homogenization of crop diversity (24), attaining the benefits of nutritional stability will require further understanding the topology of crop-nutrient networks.

These results have many important implications for our understanding of nutritional outcomes and their relationship to crop diversity. First, while our work reaffirms the existing body of research demonstrating that crop diversity is important for agricultural sustainability, it does so at a national scale. Much of the existing work in understanding crop diversity and its links to nutritional outcomes are at a field or landscape level (10). Our work answers recent calls (9) to explore crop diversity and nutritional outcomes at a larger scale through a country level analysis and incorporates both production and imports, the latter of which has been significantly important for driving an increase in the types of crops available in a given country over time.

Second, our work adds novelty by establishing a functional relationship between crop diversity and nutritional stability, and suggests that this non-linear relationship has important implications for thinking about the types of crops grown or imported in a given region and their relationship to food availability. A foundation shared by ecology and nutrition is that diversity can improve long-term functioning of complex biological systems (25, 26). Like other ecological diversity-resilience relationships, we observe that diversity loss can result in rapid non-monotonic loss of function (27). In countries where diversity is already low, our results indicate that crop failures, either through production failure or an inability to import such crops, could lead to rapid reductions in nutrient availability within a country. Moreover, multiple failures of highly important regional crops, as might occur during a drought or other extreme event, could have catastrophic nutritional impact. Such countries are thus vulnerable to a variety of potential global challenges both ecological (e.g. climate change) and economic (e.g. trade wars).

Third, that nutritional stability is stagnant or decreased over time in all regions but Asia highlights that increasing crop diversity – at least at the national level – does not necessarily lead to better nutritional outcomes. Instead, the wide variability in nutritional stability across countries, despite the overall increase in crop diversity, highlights clear vulnerabilities both across regions and within regions. Africa has the greatest inter-regional variability, demonstrating that in some cases neighboring countries have very different capacities and stabilities of crop nutrients in their food supply chain in any given year. This variability is likely driven by multiple factors including the capacity of a country to trade (28), and in country food availability as a result of war or political/social unrest (29–31), or climate induced disasters (32).

Finally, the important role of imports in our findings for many regions highlights that nutritional stability is market exposed. While trade can positively affect food security (33), it can also hinder nutrition efforts (34), and could be a vulnerability if imports comprise a significant portion of nutrients for a given population. Countries with a high reliance on imports are thus subject to trade wars, market shifts, and price shocks that can occur for a variety of reasons. Such countries may therefore be more likely to experience increased variability in the future, especially as climate change is expected to affect both agricultural production and markets and trade (35).

The use of these results could help inform high-level discussions within countries and regions about the key crops for a given place, and their availability via import versus domestic production. Scenario development using our metric could help identify country-specific crop additions that would maximize nutritional stability improvements. Our approach could be used to identify potential tradeoffs in production and import outcomes, at least as it relates to the availability of a given amount of nutrients in a certain place. Such potential applications also highlight the importance of identifying several caveats and important limitations. First, while we are addressing the nutrients available in a given country in a given time, we are not equating this with food security. This “availability” is only one component of food security, with access, utilization, and stability being other critical pillars (FAO 2008). Thus, even though nutritional stability is generally high in most regions and remained stagnant (or increased in Asia), this does not mean that people are not food insecure. Adequate food and nutritional security comprises much more than the factors captured in our analysis, which provides a relative measure of nutrient availability not an absolute metric of adequacy. Furthermore, there are recognized shortcomings with the existing FAO data, especially in many low-income countries (36) and we recognize that data quality is a challenge with this dataset in some cases. Nevertheless, to our knowledge, it is the best available data of its kind and scale available, so we utilize it knowing there are many opportunities to improve this work moving forward.

Despite these caveats, this work advances a novel method to assess the relationship between crop diversity and nutritional stability globally over the past fifty years. Future research could expand this work in multiple ways by combining crop nutrient availability data with nutritional intake data to better assess whether available nutrients in the supply chain are making their way into household consumption. This would more completely link crop diversity with food and nutritional security outcomes, rather than just food availability as this work as done. Furthermore, our novel network method could be advanced by exploring the importance of certain crops for a given country or region. Finally, with climate change expected to affect the yields of many globally important crops (37) and potentially cause multiple crop failures at once (32), this type of analysis could advance our understanding of food system vulnerability to specific crop failures, and provide guidance on climate adaptation efforts or crop diversification strategies to safeguard against climate change. Resilience is now a central paradigm in many sectors – humanitarian aid, disaster risk reduction, climate change adaptation, social protection. Erosion of biological diversity typically leads to loss of ecosystem functioning and services, likewise loss of crop diversity may to lead to potentially drastic shifts in nutritional stability. Together this, and future analyses, have the potential to direct the protection or restoration of crop diversity so as to best support nutrient availability that is stable given current and future challenges.

## Materials and Methods

### Data sources

Crop composition information came from annual national-level food balance data for years 1961 to 2016 obtained from the FAOSTAT database (Food and Agriculture Organization of the United Nations; http://fao-stat.fao.org/) for 201 nations that comprise >95% of the global population. We focus our analysis on yield of crop plants (hg/ha) available for human consumption. To explore the impact of international trade on nutrient stability, we created two food supply scenarios by evaluating national yield separately as its production (P) and production plus imports (PI) components. We linked annual national crop lists for each scenario with crop-specific nutrient data reported in the Global Expanded Nutrient Supply (GENuS) database (Smith et al 2016), a global dataset of nutrient supply of reports the amount of nutrient available per 100g of edible food for 23 nutrients across 225 food categories. We focused on 17 nutrients (calcium, carbohydrates, copper, fiber, folate, iron, magnesium, niacin, phosphorous, potassium, protein, riboflavin, thiamin, vitamin A, vitamin B6, vitamin C, and Zinc), omitting calories, fats, water, ash and refuse. Linking these two datasets results in 22,400 crop-nutrient networks, one for each country, year, and food supply scenario. However, the nutritional contribution of some foods is often negligible, yet because links in our bipartite crop-nutrient networks are treated as binary these would still be considered as contributing to stability. To overcome this minimum quantities problem (38), we calculated the proportion daily value per serving (PDV) for each country for each food as: PDV = 100*N*((Pop/Prod)/DV), where N is the grams of nutrient per 100 grams of food (as provided by GENuS), Pop is a country’s population (United Nations, 2019), Prod is the amount of that food produced (FAOSTAT, 2019) and DV is the daily value of the nutrient in grams (FDA, 2019). We used PDVs as link weights in each network and then define a threshold on the computed link weight, below which we consider the link as nonexistent. To do this we compute the stability for each country in 2016 at values of the threshold cutoff ranging from 0 to 0.7. The mean of the stabilities decreases monotonically as a function of this cutoff, but the variance shows a plateau from 0 to 0.1 before decreasing. In order to magnify between-country variation while removing negligible links, we choose the threshold of 0.1, at the upper end of this plateau. Data availability varied among global regions and missing data values were more common in the early years of the data set, we therefore removed countries that did not have at least 20 years reported (N = 17 countries), leaving 184 countries and 19,044 networks.

### Formalizing nutritional stability (R_N_)

Our algorithm processes networks through permutation of crop species removal order to generate an attack tolerance curve and uses the area under this curve as a coefficient of stability. Consider the set of *M* crops *C = {c*_*1*_, *…, c*_*M*_*}* and the set of *P* nutrients *N = {n*_*1*_, *…, n*_*P*_*}*. We begin by constructing a bipartite graph, *G*, with left nodes consisting of the set *C* and right nodes consisting of the set *N*. Place an edge between *c*_*i*_ (the *i*-th crop) and *n*_*j*_ (the *j*-th nutrient) if crop *c*_*i*_ contains nutrient *n*_*j*_. Given a subset of crops *D* ⊂ *C*, define *nbr*(*D*) to be the neighborhood of *D*, meaning the subset of *N* that is connected by an edge to an element of *D*. Finally, given a set *A*, let |*A*| be the number of elements in *A* (thus |*N*(*D*)| is the number of elements that D is connected to). Now consider the following sampling procedure

1. Initialize *C*^*0*^ = *C*
2. At step *k*, remove one randomly selected crop from *C*^*k−1*^, leaving the set *C*^*k*^, which contains *M − k* crops. Let *X* (*k*) = |*nbr*(*C*^*k*^)|, the number of nutrients connected to *C*^*k*^.
3. Repeat Step 2 *M* times (i.e., until there are no more crops left).

Thus, at each iteration the procedure described above removes a randomly selected crop and all of its edges from *G*, and counts the number of nutrients that remain connected to rest of the graph. The procedure yields a trajectory *X*(*k*), with *k* ranging from 1 to *M*, which we will call a sample. Repeating this procedure multiple times yields multiple samples *X*_*i*_(*k*). We generate an average attack tolerance and take its integral as our coefficient of nutritional stability (*R*_*N*_). This unitless measure represents how robust a food system is to the sequential elimination of crops.

### Statistical analyses

To explore the relationship between crop diversity and nutritional stability (Fig. 2) we compared three non-linear functional forms (saturating, logarithmic, exponential) using non-linear mixed effects models with crop diversity as a fixed effect and country as a random effect. A saturating function (α * x/(β + x)) was selected based on AIC (Table S5) and used for subsequent analyses. We used separate non-linear models for each region to explore the relationship between nutritional stability and crop diversity averaged across years for each country. We test whether response curves for this relationship are different between regions by extracting individual country regression coefficients for the model parameters (asymptote, α; doubling time, β) from non-linear mixed effects models with country as a random effect. We then used linear models to compare parameter coefficient values between regions (Fig S1; Table S6). We conducted these analyses using production scenario crop information, but results did not qualitatively change if imports were included. To explore changes in crop diversity, nutritional stability and average crop degree over time (Fig. 3) we used separate linear mixed effects models for each region with an interaction between supply scenario and year as fixed effects, a random intercept for scenario nested in country, and a first order autoregressive (AR-1) correlation structure to account for the temporal dependency structure of our data. Finally, to explore how nutritional stability differs between macroeconomic factors (Fig. 4; small island state status, development status) we averaged crop diversity across years for each country and used linear mixed effects models with an interaction between macroeconomic status and scenario as fixed effects and country as a random effect. We performed all analyses in R (v 3.6.1) using ‘nlme’ (39) and ‘lme4’ (40) for statistical modeling.

## Acknowledgments

We are thankful to Taylor Ricketts and Molly Brown for providing thoughtful comments and feedback on earlier versions of the manuscript. We would like to thank Rishidev Chaudhuri for comments and helping formalize our nutritional network approach. Funding for this work was provided by the USDA Hatch Program (grant number VT-H02303) given to MTN. The authors declare no conflicts of interest.

## Data Accessibility

The data presented in this manuscript will be made publicly available through Figshare (doi: xx.xxxx/xxxx [to be completed before final submission]).

## SUPPLEMENTARY MATERIALS

**Table S1.** Parameter estimates testing for a non-linear relationship between nutritional stability and crops diversity. Curves fit with a saturating function (a * x/(b + x)). This functional form was selected after multiple model comparison (**Table S5**). Individual models were fit for each region. For details on regional differences see **Figure S1** and **Table S6**. Values are model coefficients with standard error in parentheses.

| Nutritional stability $\sim a * x / (b + x)$ | | | | | |
| --- | --- | --- | --- | --- | --- |
|  | Africa | Americas | Asia | Europe | Oceania |
| a | 1.097***<br>(0.042) | 1.335***<br>(0.099) | 1.131***<br>(0.052) | 0.995***<br>(0.015) | 0.979***<br>(0.027) |
| b | 4.965***<br>(0.814) | 12.574***<br>(2.486) | 6.738***<br>(1.085) | 2.353***<br>(0.366) | 2.263***<br>(0.299) |
| Observations | 50 | 39 | 46 | 33 | 15 |
| Log Likelihood | 67.642 | 39.932 | 56.713 | 71.664 | 26.597 |
| Note: . p<0.1; * p<0.05; ** p<0.01; *** p<0.001 |  |  |  |  |  |

**Table S2.** Crop diversity trends over time. Results are from region-specific linear mixed effects model with an interaction between scenario and year as fixed-effects, country nested in scenario as random effects and an autoregressive correlation structure (i.e., time-lag correlation) to account for temporal autocorrelation. Values are model coefficients with standard error in parentheses.

|  | Crop diversity |  |  |  |  |
| --- | --- | --- | --- | --- | --- |
|  | Africa | Americas | Asia | Europe | Oceania |
| Scenario | -59.466<sup>*</sup><br>(23.739) | -75.863<sup>*</sup><br>(33.737) | -130.076<sup>*</sup><br>(50.347) | -219.441<sup>***</sup><br>(36.733) | -31.948<br>(35.314) |
| Year | 0.047<sup>***</sup><br>(0.008) | 0.060<sup>***</sup><br>(0.012) | 0.071<sup>***</sup><br>(0.018) | 0.047<sup>***</sup><br>(0.013) | 0.018<br>(0.013) |
| Scenario × Year | 0.030<sup>*</sup><br>(0.012) | 0.039<sup>*</sup><br>(0.017) | 0.067<sup>**</sup><br>(0.025) | 0.113<sup>***</sup><br>(0.018) | 0.017<br>(0.018) |
| Observations | 5,546 | 4,217 | 4,577 | 3,076 | 1,628 |
| Log Likelihood | -8,590.126 | -7,130.420 | -8,581.432 | -5,601.669 | -2,366.630 |
| *Note:* . p<0.1; * p<0.05; ** p<0.01; *** p<0.001 |  |  |  |  |  |

**Table S3.** Macroeconomic factors drive nutritional stability differences. Differences (estimate standard error in parentheses) in nutritional stability between (a) developing and non-developing countries and (b) small island developing states (SIDS). Results are from separate linear mixed effects models with an interaction between macroeconomic status and scenario as fixed-effects and country as a random effect.

**(a)**
| scenario | contrast | estimate | SE | df | *t* ratio | *p* value |
| --- | --- | --- | --- | --- | --- | --- |
| P | Developing - Non-developing | -0.0674 | 0.0265 | 363 | -2.541 | 0.0115 |
| PI | Developing - Non-developing | -0.0693 | 0.0265 | 363 | -2.612 | 0.0094 |

**(b)**
| scenario | contrast | estimate | SE | df | *t* ratio | *p* value |
| --- | --- | --- | --- | --- | --- | --- |
| P | Non-SIDS - SIDS | 0.133 | 0.0257 | 363 | 5.172 | < 0.0001 |
| PI | Non-SIDS - SIDS | 0.129 | 0.0255 | 363 | 5.083 | < 0.0001 |

**Table S4.** Crop degree trends over time. Results are from region-specific linear mixed effects model with an interaction between scenario and year as fixed-effects, country nested in scenario as random effects and an autoregressive correlation structure (i.e., time-lag correlation) to account for temporal autocorrelation. Values are model coefficients with standard error in parentheses.

|  | Crop degree |  |  |  |  |
| --- | --- | --- | --- | --- | --- |
|  | Africa | Americas | Asia | Europe | Oceania |
| Scenario | -4.168<br>(7.645) | -8.645<br>(9.645) | -0.841<br>(10.455) | 35.491***<br>(5.733) | -9.692<br>(16.069) |
| Year | -0.022***<br>(0.003) | -0.024***<br>(0.003) | -0.020***<br>(0.004) | -0.011***<br>(0.002) | -0.015**<br>(0.006) |
| Scenario × Year | 0.002<br>(0.004) | 0.004<br>(0.005) | 0.0005<br>(0.005) | -0.018***<br>(0.003) | 0.005<br>(0.008) |
| Observations | 5,546 | 4,217 | 4,577 | 3,076 | 1,628 |
| Log Likelihood | -4,488.510 | -2,567.151 | -3,741.008 | -2,694.267 | -1,716.429 |
| Note: | . p<0.1; * p<0.05; ** p<0.01; *** p<0.001 |  |  |  |  |

**Table S5.** Comparing the relationship between nutritional stability and crop diversity using three saturating model forms. Based on AIC scores the saturating function α * x/(β + x) was used in subsequent analyses (Fig. 2; Table S1 & Table S6). Values are model coefficients with standard error in parentheses.

|  | Nutritional stability |  |  |
| --- | --- | --- | --- |
| | $\alpha + \beta * \log(x)$ | $\alpha * x / (\beta + x)$ | $\alpha * \exp(\beta * x)$ |
| b | 0.161***<br>(0.004) | 5.126***<br>(0.462) | 0.017***<br>(0.001) |
| a | 0.234***<br>(0.002) | 1.085***<br>(0.023) | 0.592***<br>(0.017) |
| Observations | 183 | 183 | 183 |
| Log Likelihood | 184.529 | 200.093 | 125.840 |
| AIC | -359.059 | -390.186 | -241.679 |
| Note: | . p<0.1; * p<0.05; ** p<0.01; *** p<0.001 |  |  |

**Table S6.** Comparing differences in parameter estimates for region-specific relationship between nutrient stability and crop diversity (Africa is contrast reference). Curves fit with a saturating function (α * x/(β + x)) via non-linear mixed effects models (see Methods) and coefficient values were extracted from random effects for each country. Values are model coefficients with standard error in parentheses.

|  | Saturating function parameter |  |
| --- | --- | --- |
| | $\alpha$ | $\beta$ |
| Americas | 0.238***<br>(0.00000) | 7.619***<br>(0.100) |
| Asia | 0.034***<br>(0.00000) | 1.762***<br>(0.095) |
| Europe | -0.102***<br>(0.00000) | -2.602***<br>(0.105) |
| Oceania | -0.118***<br>(0.00000) | -2.692***<br>(0.137) |
| Observations | 183 | 183 |
| *Note:* . p<0.1; * p<0.05; ** p<0.01; *** p<0.001 |  |  |

**Figure S1.**
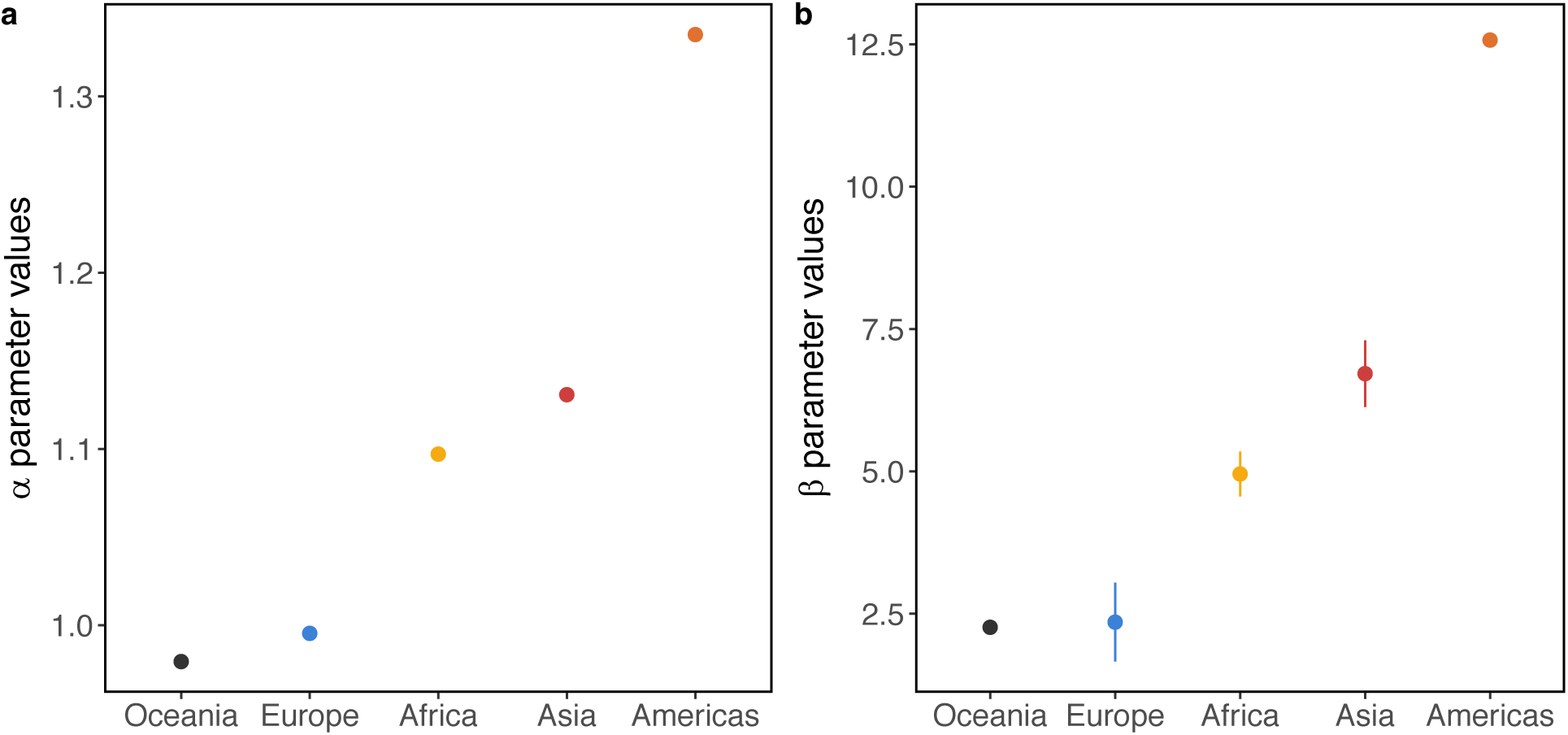
Slope parameter estimates from non-linear mixed effects models relating crop diversity and nutritional stability (see Fig. 1). Curves fit with a saturating function (α * x/(β + x)) via non-linear mixed effects models (**Table S6**; see Methods) and coefficient values were extracted from random effects for each country. Points depict the average ± sd across countries for the α parameter (a) and β parameter (b).

**Figure S2.**
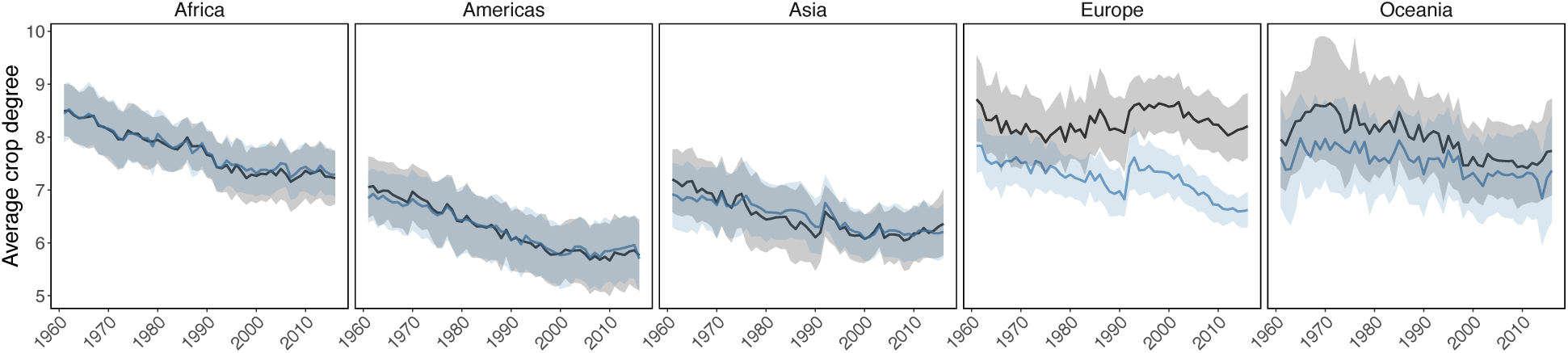
Average degree of crops in crop-nutrient networks decreased over time. Only Europe exhibited scenario-dependent differences, with production plus imports (blue) decreasing more than production alone (black), see **Table S4** for statistics. Trend lines depict means ± 95% confidence intervals.

